# Targeting the 3’ splice site by a decoy oligonucleotide attenuates U2AF1 splicing activity and inhibits leukemia

**DOI:** 10.64898/2026.06.30.735457

**Authors:** Carla Azar-Koussa, Marina Sakran, Efraim Rahamim, Anurag V Prabhu, Samar Salem, Adina Heinberg, Zahava Siegfried, Meital Ben-David-Naim, Eran Zimran, Erez Y. Levanon, Zvi Granot, Rotem Karni

## Abstract

Mutations in spliceosomal genes are a hallmark of myeloid malignancies, yet leukemic cells remain dependent on residual splicing activity, exposing a therapeutic vulnerability. Here, we present an RNA decoy strategy that disrupts 3′ splice site recognition by competitively engaging spliceosomal components. We engineered a chemically stabilized RNA decoy that mimics the 3′ splice site (3′SS decoy), sequestering proteins involved in 3′ splice site recognition from endogenous pre-mRNA targets. Although the decoy is expected to engage multiple components the 3’ splice site recognition complex, U2AF1 served as the primary molecular readout to assess target engagement and downstream effects. Lipid nanoparticle (LNP) encapsulation enabled efficient intracellular delivery into leukemic cell lines and patient-derived blasts, where the decoy directly engaged the splicing machinery, induced widespread splicing alterations, impaired leukemic cell fitness in vitro, and significantly reduced leukemia burden in vivo. These effects were independent of spliceosomal mutational status, establishing decoy-mediated disruption of splicing factor activity as a mechanistically targeted therapeutic strategy for myeloid malignancies.

## Introduction

Splicing is a fundamental step in gene expression that enables the generation of multiple protein isoforms from a single gene. This process is frequently dysregulated in human disease, including cancer, where splicing factors are often overexpressed and can contribute to malignant transformation when aberrantly regulated^1^. In hematological malignancies, recurrent mutations in core spliceosomal components have emerged as a defining feature, with *U2AF1*, *SF3B1*, *SRSF2*, and *ZRSR2* among the most frequently affected genes^2^. These mutations are typically heterozygous, mutually exclusive missense alterations targeting highly conserved residues^3^, highlighting a selective pressure to preserve partial spliceosomal function while altering its activity. Splicing factor mutations are observed in approximately 20-30% of de novo acute myeloid leukemia (AML) cases and in more than 50% of patients with myelodysplastic syndromes (MDS) and secondary AML, underscoring their central role in disease pathogenesis^4^. Both MDS and AML are clonal disorders of hematopoietic stem and progenitor cells, characterized by ineffective hematopoiesis, bone marrow failure, and peripheral cytopenias, together with the accumulation of immature and functionally impaired myeloid cells^5^. Collectively, these findings highlight a critical dependency of malignant myeloid cells on a dysregulated, yet essential, splicing machinery.

Importantly, this dependency is not restricted to cells harboring spliceosomal mutations. The predominance of heterozygous mutations and their mutual exclusivity suggest that loss of splicing factor function is not tolerated, indicating that leukemic cells remain reliant on residual splicing activity for survival. This raises the possibility that perturbing splicing factor function may represent a broader therapeutic vulnerability, extending beyond genetically defined subsets to leukemic cells that retain wild-type spliceosomal components. For example, a previously developed therapeutic, Pladienolide B, is a potent cancer cell growth inhibitor targeting the SF3B1 subunit of the spliceosome^6,7^. Accordingly, targeting splicing factor activity may provide a strategy to disrupt this essential dependency across a wider spectrum of myeloid malignancies.

Building on this concept, we sought to therapeutically exploit the dependence of leukemic cells on functional splicing machinery. Here, we present a decoy oligonucleotide-based strategy^6,7^ to selectively disrupt splicing factor activity at the 3′ splice site. This approach employs engineered RNA decoys that mimic the consensus 3′ splice site motif, thereby competitively engaging components of the spliceosomal machinery. During the early stages of major spliceosome assembly, key cis-regulatory RNA elements including the 5′ splice site, branch point sequence, polypyrimidine tract, and 3′ splice site are recognized by distinct spliceosomal components. Specifically, U1 small nuclear ribonucleoprotein (snRNP) binds the 5′ splice site, SF3B1 recognizes the branch point sequence, and the U2AF2-U2AF1 heterodimer engages the polypyrimidine tract and 3′ splice site recruiting the U2 snRNP^8^. Given that U2AF1 specifically recognizes the AG dinucleotide at the 3′ splice site downstream of the polypyrimidine tract^9,10^ we hypothesized that a synthetic RNA decoy incorporating this recognition motif could competitively inhibit U2AF1 function by sequestering the factor, as well as additional 3’ splice site factors, away from endogenous pre-mRNA substrates, thereby impairing splice site recognition and downstream splicing activity. While the decoy is designed to target the 3′ splice site recognition complex in a general manner, we use U2AF1 as a primary molecular readout to characterize binding and downstream splicing effects. Since transcription and replication are hyper activated in cancer cells, efficient splicing is required to enable proper gene expression and cell survival^11,12^. By perturbing the interactions of the 3’ splice site with its binding proteins and disrupting splicing fidelity, we expect to attenuate the survival of leukemic cells, which are highly dependent on splicing fidelity compared to normal cells.

In addition to targeting the therapeutic potential of splicing factors in myeloid neoplasms, it is crucial to address the major challenge of effectively delivering RNA-based therapies into cancer cells. Although there have been substantial advances in the area of oligonucleotide therapeutic research in the last decade^13^ the delivery of RNA-based therapeutics to many cells and tissues continues to be a major obstacle. One way to overcome the challenge of RNA stability and delivery is through the modification of the RNA-based oligonucleotides^14^. Additionally, a more recent approach is the use of nanoparticles, highly customizable delivery system in terms of size, chemical composition and surface chemistry. Lipid nanoparticles (LNPs) are versatile lipid-based carriers designed to encapsulate various RNA payloads, ensuring their delivery to cells while protecting them from degradation and preventing the activation of RNA-sensing mechanisms and subsequent innate immune responses^15^. The decoy oligonucleotides were encapsulated in LNPs to enable efficient delivery into leukemic cells.

To evaluate this approach, we first assessed the ability of the engineered RNA decoy to engage components of the 3′ splice site recognition complex, using U2AF1 as a primary molecular readout. We then examined the downstream impact of decoy-mediated sequestration on global splicing patterns in leukemic cells. Finally, we investigated the functional consequences of this perturbation on leukemic cell survival both in vitro and in vivo, establishing the therapeutic potential of targeting spliceosomal dependencies through LNP- encapsulated decoy-based strategies.

## Results

### Design of the 3’SS decoy oligonucleotide

Building on our prior development of decoy oligonucleotides targeting the splicing factors RBFOX2, PTBP1, and SRSF1^16^, we sought to extend this strategy to target the 3′ splice site by designing a decoy that perturbs a central node of spliceosome recognition, thereby exposing a potential vulnerability of leukemic cells to splicing disruption^17,18^. In order to directly inhibit U2AF1-RNA interactions, we engineered a chemically stabilized synthetic RNA decoy oligonucleotide that mimics the canonical 3′ splice site recognition motif, with the goal of competitively sequestering U2AF1 from endogenous pre-mRNA targets. The decoy consists of a 39-nucleotide single-stranded RNA containing three tandem repeats of a consensus U2AF1-binding motif, UUUUUUUUAGCCC (**Fig. 1a**). To enhance nuclease resistance, cellular uptake, and intracellular stability, the oligonucleotide was chemically modified with 2′-O-methyl (2′-OMe) substitutions across the central 31 nucleotides, flanked by 2′-O-methoxyethyl (2′-MOE) modifications at the terminal four nucleotides on each end. A phosphorothioate (PS) backbone was incorporated throughout the sequence. For selected in vitro applications, oligonucleotides were additionally labeled at the 5′ end with either Cy3 or biotin. A scrambled oligonucleotide of identical chemistry served as a control in all experiments.

**Figure 1.**
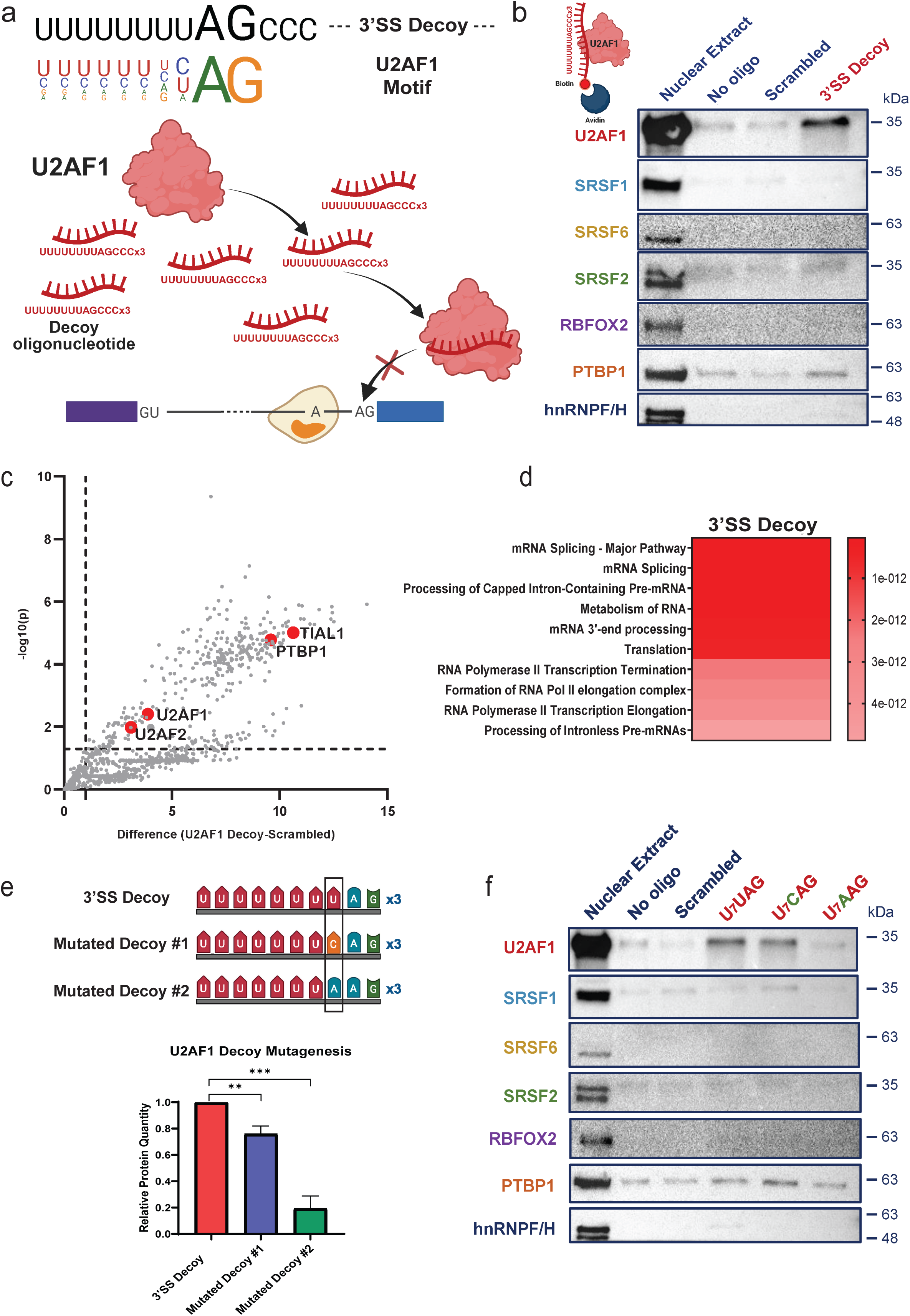
Characterization and validation of the 3’SS decoy oligonucleotide. (a) Alignment of the U2AF1 consensus RNA-binding motif (colored nucleotides) with the designed decoy oligonucleotide sequence (black nucleotides) (top). Schematic illustration of U2AF1 inhibition through competitive binding to the decoy oligonucleotide (bottom). (b) Western blot analysis of proteins pulled-down using biotin-conjugated 3’SS decoy or scrambled control oligonucleotide from K562 nuclear extracts. (c) Scatter plot showing significantly enriched proteins (above the dashed significance threshold) in pull-downs performed with the biotin-conjugated 3’SS decoy compared to the scrambled control, followed by mass spectrometry analysis. Known splicing factors: U2AF2, U2AF1, PTBP1, TIAL1 are marked in red. (d) Reactome pathway enrichment analysis of significantly enriched proteins identified in the 3’SS decoy pull-down followed by mass spectrometry. Pathways were filtered using a significance threshold of *P* < 0.05 and FDR < 0.05; the top 10 pathways are shown. (e) Schematic representation of the 3’SS decoy sequence (first row) and two mutant variants, each containing a single nucleotide substitution at one position (top). The mutation site is marked with a back rectangular box. Quantification of Western blot (bottom). Unpaired two-tailed Student’s *t*-test was performed on data. Data is shown as mean ± s.d. P-values: ** p < 0.01, *** p < 0.001. (f) Western blot analysis of proteins pulled down using biotin-conjugated 3’SS decoy, mutant decoys, or scrambled control oligonucleotide from K562 nuclear extracts. In all panels, *n* ≥ 3 independent experiments.

### The 3’SS decoy selectively binds U2AF1 and associated splicing factors

Having established the design of a 3′ splice site decoy, we next sought to determine whether it physically engages U2AF1 and associated spliceosomal components in a sequence-specific manner. To this end, biotin-mediated pull-down assays using nuclear extracts from K562 cells were performed. These pull-down experiments demonstrated robust enrichment of U2AF1 pulled-down by the 3’SS decoy, but not by the scrambled control (**Fig. 1b; Supplementary Fig. 1a**). Proteomic analysis of decoy-associated complexes by mass spectrometry confirmed significant enrichment of U2AF1 and additional splicing factors, consistent with the recruitment of spliceosomal assemblies. Notably, the decoy also bound the RNA binding proteins PTBP1 and TIAL1, likely reflecting the presence of a polypyrimidine tract within the decoy sequence (**Fig. 1c; Supplementary Fig. 1b**). A STRING analysis of the functional protein association networks of the significantly enriched proteins revealed clustering into transcription, translation, and mRNA processing modules, reflecting the functional coupling of these pathways and the co-purification of interacting protein complexes with the decoy during pull-down (**Supplementary Fig. 1c**). Reactome pathway and gene ontology enrichment analyses of the mass-spectrometry data identified enrichment of proteins involved in mRNA splicing pathways, RNA processing, mRNA metabolic processing, and ribonucleoprotein complex assembly (**Fig. 1d, Supplementary Fig. 1d**).

To assess sequence specificity, we introduced point mutations at the nucleotide positioned three bases upstream of the 3′ splice junction within the decoy, a critical determinant of U2AF1 recognition. Biotin-mediated pull-down assays using these mutated decoys revealed that substitution of uridine with cytidine reduced binding to U2AF1 significantly, whereas substitution with adenosine nearly abolished interaction (**Fig. 1e, f**). Together, these findings demonstrate that the 3’SS decoy selectively and sequence-dependently engages U2AF1 and its interacting splicing machinery, establishing a functional platform to interrogate the consequences of sequestration of 3′ splice site binding factors in leukemic cells.

### The 3’SS decoy induces widespread alterations in alternative splicing

Since U2AF1 is an important component of the spliceosome complex required for pre-mRNA splicing, we next investigated whether decoy-mediated 3’SS/U2AF1 sequestration leads to transcriptome-wide splicing perturbations. RNA- seq analysis of K562 cells treated with LNP-delivered 3’SS decoy or scrambled oligonucleotides revealed distinct transcriptional and splicing profiles. Principal component analysis and hierarchical clustering resulted in a clear separation between treatment groups (**Fig. 2a, b**). Analysis using PSI-Sigma identified 566 genes with significant splicing changes (ΔPSI > 10%) upon 3’SS decoy treatment when compared to scrambled oligonucleotide treatment (**Fig. 2c).** In parallel, differential gene expression analysis also revealed numerous transcriptional alterations (**Supplementary Fig. 2a-c**). Splicing changes in selected targets, in addition to previously reported U2AF1-regulated transcripts^19^, were validated by RT-PCR (**Fig. 2d; Supplementary Fig. 2d**), with K562 cells knocked-down for U2AF1 serving as a positive control (**Fig. 2d; Supplementary Fig. 2e, f**). Motif enrichment analysis (XSTREME) of U2AF1 regulated splice sites revealed significant enrichment of UNAG and U_7_AG motifs, consistent with canonical U2AF1 binding preferences (**Fig. 2e**). Reactome pathway analysis^20^ further indicated that affected genes are enriched in cancer-relevant signaling pathways (**Fig. 2f**). Taken together, these data reveal that targeting the 3′ splice site with a synthetic decoy is sufficient to alter the leukemic transcriptome through widespread disruption of U2AF1-dependent splicing.

**Figure 2.**
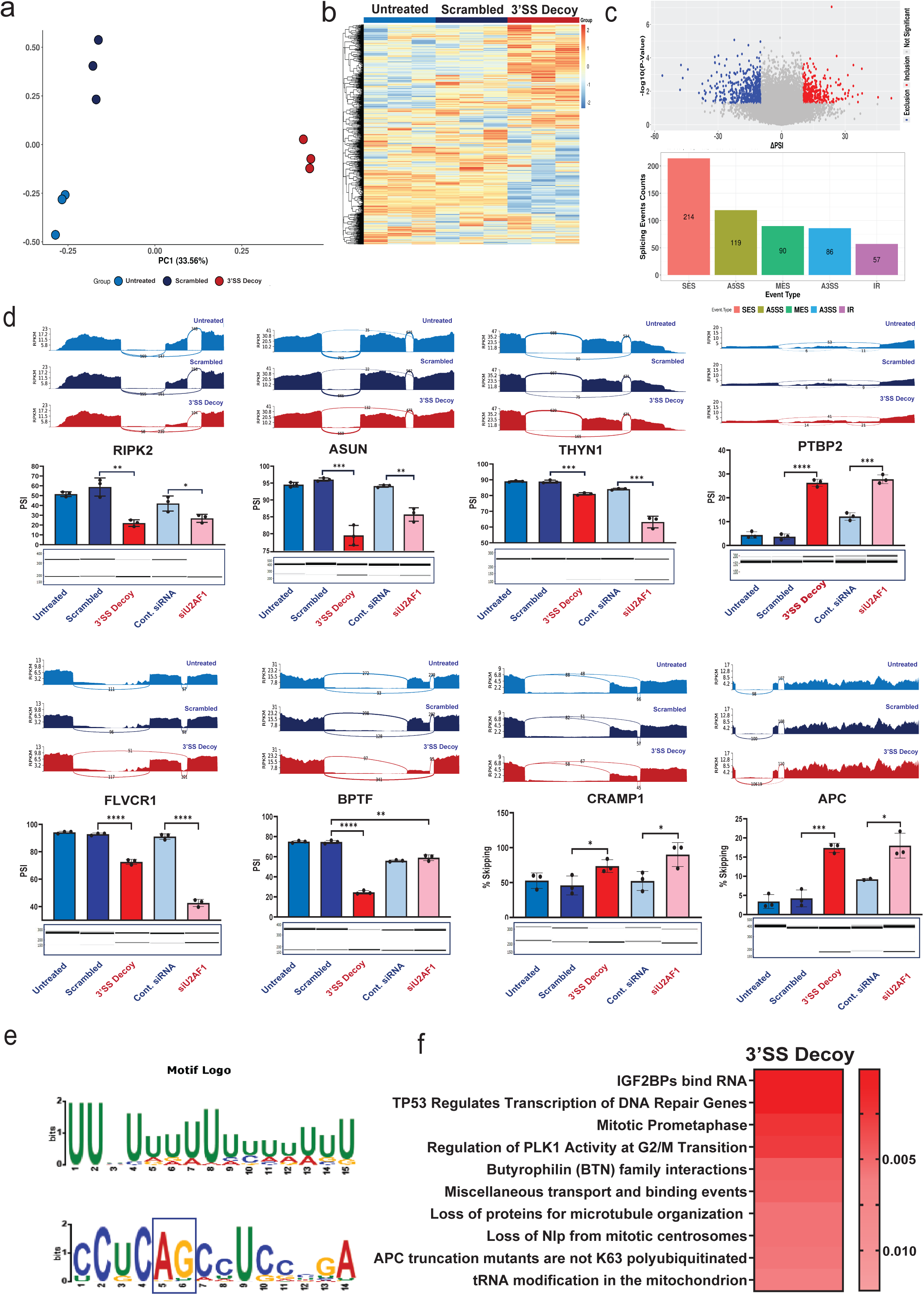
The 3’SS decoy induces leukemic cell sensitivity mediated through splicing changes. (a) PCA based on Percent Spliced In (PSI) values of detected splicing events across samples. Each point represents a sample and is colored by experimental condition: untreated (light blue), scrambled control (dark blue), and 3’SS decoy-treated cells (red). The PCA demonstrates separation of the decoy-treated samples from the control groups based on their splicing profiles. (b) Heatmap of PSI values for alternative splicing events across samples. Rows represent splicing events and columns represent individual samples. Hierarchical clustering was performed on the events, and PSI values are shown as scaled values (z-scores). Samples are annotated by treatment group as indicated. (c) Differential splicing analysis between scrambled control and 3’SS decoy-treated samples. Volcano plot showing the relationship between the magnitude of splicing change (ÄPSI) and statistical significance (−log10 p-value). Events meeting significance thresholds (|ΔPSI| > 10%, p-value < 0.05, FDR < 0.05) are highlighted, with increased exon inclusion in decoy-treated samples shown in red and increased exon exclusion shown in blue (top). Distribution of significant alternative splicing events by event type identified by PSI-Sigma, including single exon skipping (SES), alternative 5′ splice site (A5SS), multiple exon skipping (MES), alternative 3′ splice site (A3SS), and intron retention (IR) (bottom). (d) Representative sashimi plots are shown illustrating RNA-seq read coverage and splice junction support for selected alternatively spliced events identified in the analysis. For each event, the plots display read coverage across the exon and its flanking regions, along with arcs indicating splice junction reads connecting adjacent exons. Samples are shown for untreated, scrambled control, and 3’SS decoy-treated conditions. These plots illustrate the differential exon inclusion or exclusion patterns associated with 3’SS decoy treatment. Data are presented as mean ± SD from *n* = 3 independent experiments. Statistical significance was assessed using an unpaired two-tailed Student’s *t*-test. RT-PCR validation of U2AF1 targets. Primers were designed to amplify regions flanking the alternatively spliced exons (see Supplementary Table). Shown are representative LabChip electropherograms and corresponding quantification of RT-PCR products from RNA isolated from K562 cells treated with the indicated scrambled control, 3’SS decoy, siRNA control, siU2AF1, oligonucleotide- encapsulated LNPs at 500nM after 72 hours. Unpaired two-tailed Student’s *t*-test was performed on data. Data is shown as mean ± s.d. P-values: ns= no asterisk p > 0.05, * p < 0.05, ** p < 0.01, *** p < 0.001, **** p < 0.0001. (e) Motif enrichment analysis of sequences surrounding differentially spliced events. Sequence logos represent enriched RNA motifs identified within regions extending ±250 bp from the alternatively spliced sites. Motif enrichment was assessed using a biased analysis, in which known U2AF1 binding motifs were provided as input (UNAG, UUUUUUUUAG). The displayed motifs illustrate enriched sequence patterns identified in the biased analysis, with corresponding statistical significance E-values (<0.05). Letter height in the motif logos reflects the relative nucleotide frequency and information content at each position. (f) Reactome pathway analysis of the significant differentially spliced genes between scrambled and 3’SS decoy treated cells. Genes entered into the analysis had an imposed cut-off (|ΔPSI| > 10, p-value <0.05, FDR < 0.05). Pathways were filtered for a statistical threshold of *P* < 0.05; the top 10 pathways are shown. In all panels, *n* ≥ 3 independent experiments.

### Delivery of the 3’SS decoy into leukemic cells inhibits proliferation and induces apoptosis

Having demonstrated that the decoy engages U2AF1 and associated spliceosomal components, we next sought to determine whether it can be efficiently delivered into leukemic cells and elicit functional consequences. For this purpose, the decoy and scrambled oligonucleotides were encapsulated in LNPs, yielding particles with the size distribution in the range of 100nm and polydispersity index of less than 0.12 (**Supplementary Fig. 3a**). Fluorescence-based analyses demonstrated efficient uptake of the CY5-labeled LNPs and intracellular delivery of the CY3-labeled RNA cargo in K562 cells (**Fig. 3a, b, Supplementary Fig. 3b**). Interestingly, in co-culture experiments comprising of both K562 cells and primary human white blood cells, LNP uptake was preferentially observed in leukemic cells, with minimal uptake in normal cells (**Supplementary Fig. 3c, d**). Similar delivery efficiency was confirmed in primary patient-derived leukemic samples from three leukemia patients (**Supplementary Fig. 3e**). After observing an efficient delivery strategy of RNA-oligonucleotides into cell lines and patient samples, we next sought to assess the functional and biological effects of the decoy. Functionally, treatment of K562 cells with the 3’SS decoy resulted in a reduction in cell proliferation and an increase in apoptosis compared to cells treated with scrambled oligonucleotides (**Fig. 3c, d**). Collectively, these results demonstrate that LNP-mediated delivery enables efficient intracellular delivery of the decoy and induces cytotoxic effects in leukemic cells, supporting its potential as a therapeutic strategy.

**Figure 3.**
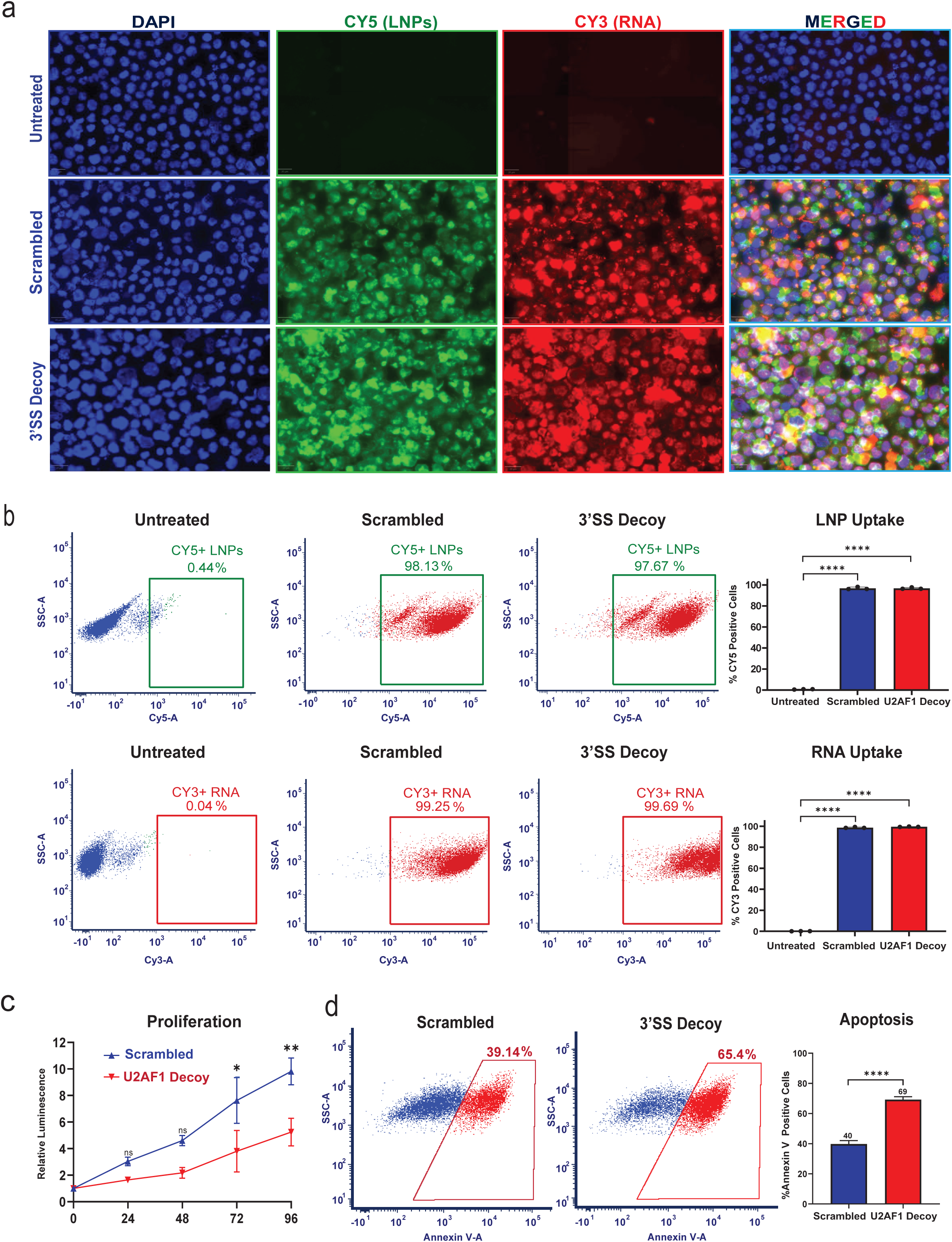
Efficient delivery and functional activity of the 3’SS decoy in leukemic cells. (a) K562 cells were incubated for 24 hours at 37°C with Cy3-labeled RNA oligonucleotides (scrambled or 3’SS decoy) encapsulated in Cy5-labeled LNPs. Representative immunofluorescence images showing cellular uptake (nuclei stained with DAPI) (bottom). (b) Representative flow cytometry plots showing uptake of Cy5-labeled LNPs encapsulating Cy3 (PE)-labeled RNA oligonucleotides (scrambled oligonucleotide or 3’SS decoy oligonucleotide) in K562 cells, 72 h after treatment with 500nM of LNP encapsulated oligonucleotides incubated at 37°C (top). Quantification of Cy5 and Cy3 (PE) positive cells. A total of 10,000 events were acquired per sample (bottom). Unpaired two-tailed Student’s *t*-test was performed on data. Data is shown as mean ± s.d. P-values: **** p < 0.0001. (c) CellTiter-Glo proliferation assay performed on K562 cells treated with 500nM LNP-encapsulated RNA (scrambled control or 3’SS decoy). Two-way ANOVA *t*-test was performed on data. Data is shown as mean ± s.d. P-values: ns: p > 0.05, * p < 0.05, ** p < 0.01. (d) Annexin V apoptosis assay in K562 cells treated with 500nM LNP-encapsulated RNA (scrambled control or 3’SS decoy). Cells were harvested after 72 hours, stained with Annexin V, and analyzed by flow cytometry (*n* = 3; 10,000 events/sample). Unpaired two-tailed Student’s *t*-test was performed on data. Data is shown as mean ± s.d. P-values: **** p < 0.0001. In all panels, *n* ≥ 3 independent experiments.

### Functional modulation of U2AF1 splicing targets, phenocopies decoy-induced effects

To determine whether specific splicing events contribute to the observed phenotype, we functionally interrogated key U2AF1 targets. Several splicing events regulated by the 3’SS decoy have a direct and indirect connection to different aspects of cancer development and progression, including *APC*^21^, *PTBP2*^22^, *FLVCR1*^23^, *RIPK2*^24^, and *BPTF*^25^ (**Fig. 2d**). Among the regulated targets, we identified Receptor interacting serine/threonine kinase 2 *(RIPK2*) and Bromodomain PHD Finger Transcription factor (*BPTF)* as alternatively spliced targets after 3’SS decoy treatment. RIPK2 is known to be involved in signaling complexes in immune pathways^24^, and BPTF is known to be the largest subunit of the nucleosome remodeling factor (NURF), playing an important role in gene regulation^25^. Both genes have established roles in oncogenesis^26–28^ and *RIPK2* splicing has previously been reported to be affected by the U2AF1 S34F point mutation in leukemia^19^. However, the functional relevance of the splice isoforms of both genes in leukemia remains unclear. To directly assess the impact of splicing modulation, we used CRISPR- Cas9 to induce exon skipping. To induce skipping of exon 2 of *RIPK2*, we designed two CRISPR sgRNAs surrounding the 5’ splice site of exon 2 (**Fig. 4a**). Both these sgRNAs triggered efficient exon 2 skipping (**Fig. 4b**). K562 cells expressing these sgRNAs had a decreased proliferation rate and increased death through apoptosis as compared to cells with control gRNAs (**Fig. 4c, d**). We similarly induced the skipping of exon 27 of *BPTF* by designing CRISPR sgRNAs around either the 3’ or 5’ splice sites of that exon (**Fig. 4e, f**). K562 cells expressing these *BPTF* sgRNAs also had a decreased proliferation rate and increased death through apoptosis (**Fig. 4g, h**). Together, these findings demonstrate that at least some of the splicing events regulated by U2AF1 and affected by the 3’SS decoy contribute directly to oncogenesis. The perturbation of individual U2AF1-regulated splicing events is sufficient to impair leukemic cell fitness, supporting a mechanistic link between decoy-induced splicing alterations and cellular phenotype.

**Figure 4:**
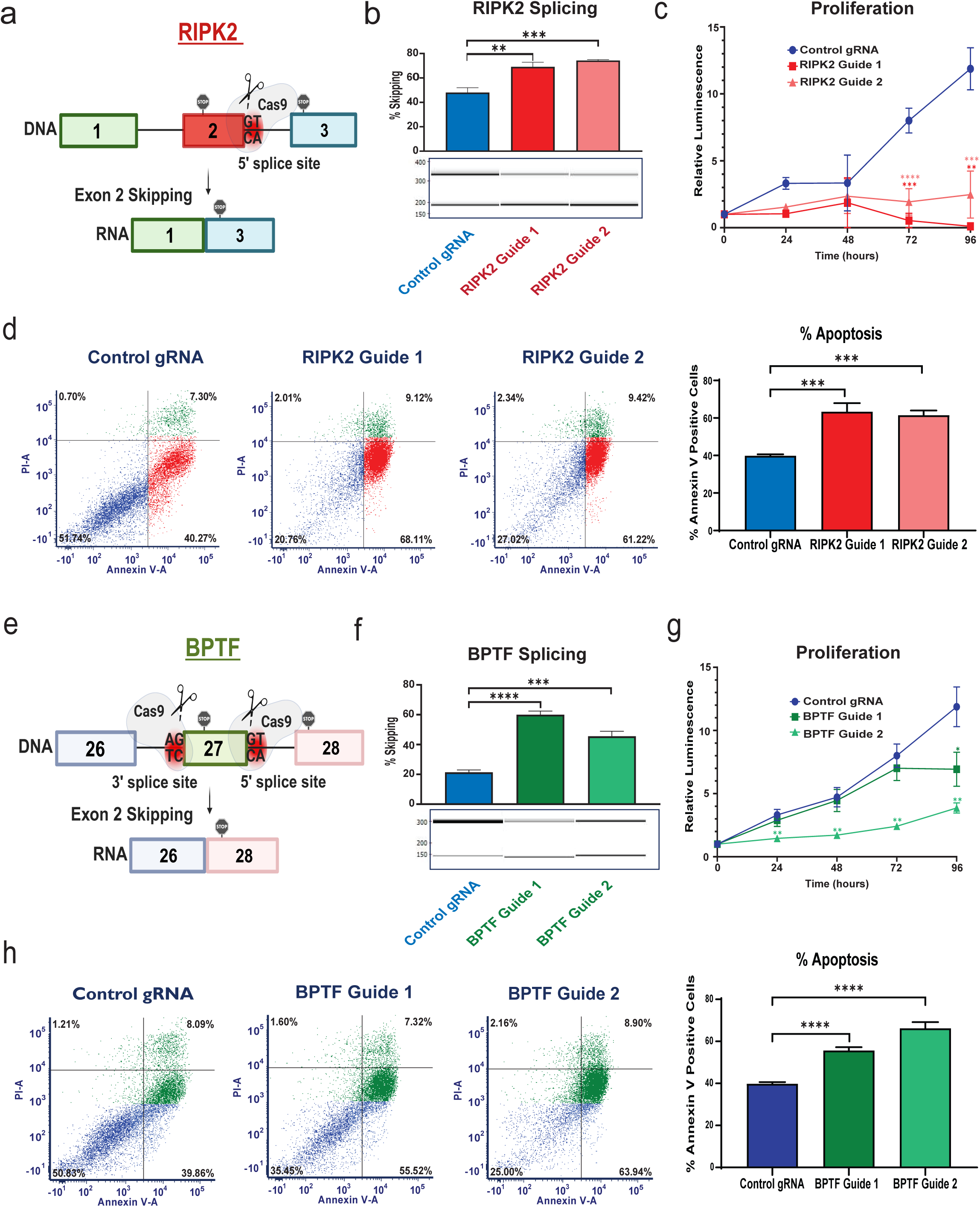
Modulation of the splicing of U2AF1 targets induces leukemic cell sensitivity. (a) Scheme of modulation of *RIPK2* exon 2 splicing using the CRISPR-Cas9 system, where guide RNAs were designed around the 5’ splice site of exon 2. (b) RT-PCR analysis of *RIPK2* splicing in K562 cells transduced with two *RIPK2* sgRNAs and a control gRNA. Shown are representative LabChip electropherograms and a corresponding quantification of RT-PCR products. Unpaired two-tailed Student’s *t*-test was performed on data. Data is shown as mean ± s.d. P-values: ** p < 0.01, *** p < 0.001. (c) CellTiter-Glo proliferation assay performed on K562 cells transduced with two *RIPK2* sgRNAs and a control gRNA. Two-way ANOVA *t*-test was performed on data. Data is shown as mean ± s.d. P-values: ns= no asterisk p > 0.05, ** p < 0.01, *** p < 0.001, **** p < 0.0001. (d) Representative flow cytometry plots of annexin V apoptosis assay in K562 cells transduced with two *RIPK2* sgRNAs and a control gRNA. A total of 10,000 events were acquired per sample. Quantification of PI negative and Pacific Blue (Annexin V) positive cells. Data is shown as mean ± s.d. P-values: *** p < 0.001. (e) Scheme of modulation of *BPTF* splicing using the CRISPR-Cas9 system, where guide RNAs were designed around the 3’ and 5’ splice site of exon 27. (f) RT-PCR analysis of *BPTF* splicing in K562 cells transduced with two *BPTF* sgRNAs and a control gRNA. Shown are representative LabChip electropherograms and corresponding quantification of RT-PCR products. Unpaired two-tailed Student’s *t*-test was performed on data. Data is shown as mean ± s.d. P-values: *** p < 0.001, **** p < 0.0001. (g) CellTiter-Glo proliferation assay performed on K562 cells transduced with two *BPTF* sgRNAs and a control gRNA. Two-way ANOVA *t*-test was performed on data. Data is shown as mean ± s.d. P-values: ns= no asterisk p > 0.05, * p < 0.05, ** p < 0.01. (h) Annexin V apoptosis assay in K562 cells transduced with two *BPTF* sgRNAs and a control gRNA. Quantification of PI negative and Pacific Blue (Annexin V) positive cells. A total of 10,000 events were acquired per sample. Unpaired two-tailed Student’s *t*-test was performed on data. Data is shown as mean ± s.d. P-values: **** p < 0.0001. In all panels, *n* ≥ 3 independent experiments.

### The 3’SS decoy suppresses leukemia progression in vivo

Finally, we evaluated whether decoy-mediated splicing disruption translates into therapeutic efficacy in vivo. NOD.Cg- *Prkdc^scid^ Il2rg ^tm1Wjl^* /SzJ (NSG) immunodeficient mice were intravenously injected with luciferase-expressing K562 cells, followed by intravenous injections of LNP- encapsulated 3’SS decoy or scrambled control oligonucleotides starting three days post-engraftment. Mice received intravenous administrations of LNP-encapsulated 3’SS decoy or scrambled control oligonucleotides (1mg/kg) every 2-3 days (**Fig. 5a; Supplementary Fig. 4a**). Longitudinal bioluminescence imaging revealed a significant reduction in leukemic burden in decoy-treated mice compared to controls, both in the whole mice and their harvested organs, in two different experiments (**Fig. 5b-e; Supplementary Fig. 4b-d**). Additionally, RNA extracted from infiltrated livers had reduced levels of human GAPDH as compared to mouse GAPDH in 3’SS- decoy treated mice, further confirming the significant reduction in leukemic burden in the mice organs (**Fig. 5f**). To confirm target engagement in vivo, RNA was extracted from infiltrated livers. RT-PCR analysis revealed significant alterations in splicing of validated human U2AF1 targets (**Fig. 5g**). Collectively, these results establish that systemic delivery of the 3’SS decoy effectively inhibits leukemia progression in vivo through modulation of alternative splicing programs.

**Figure 5:**
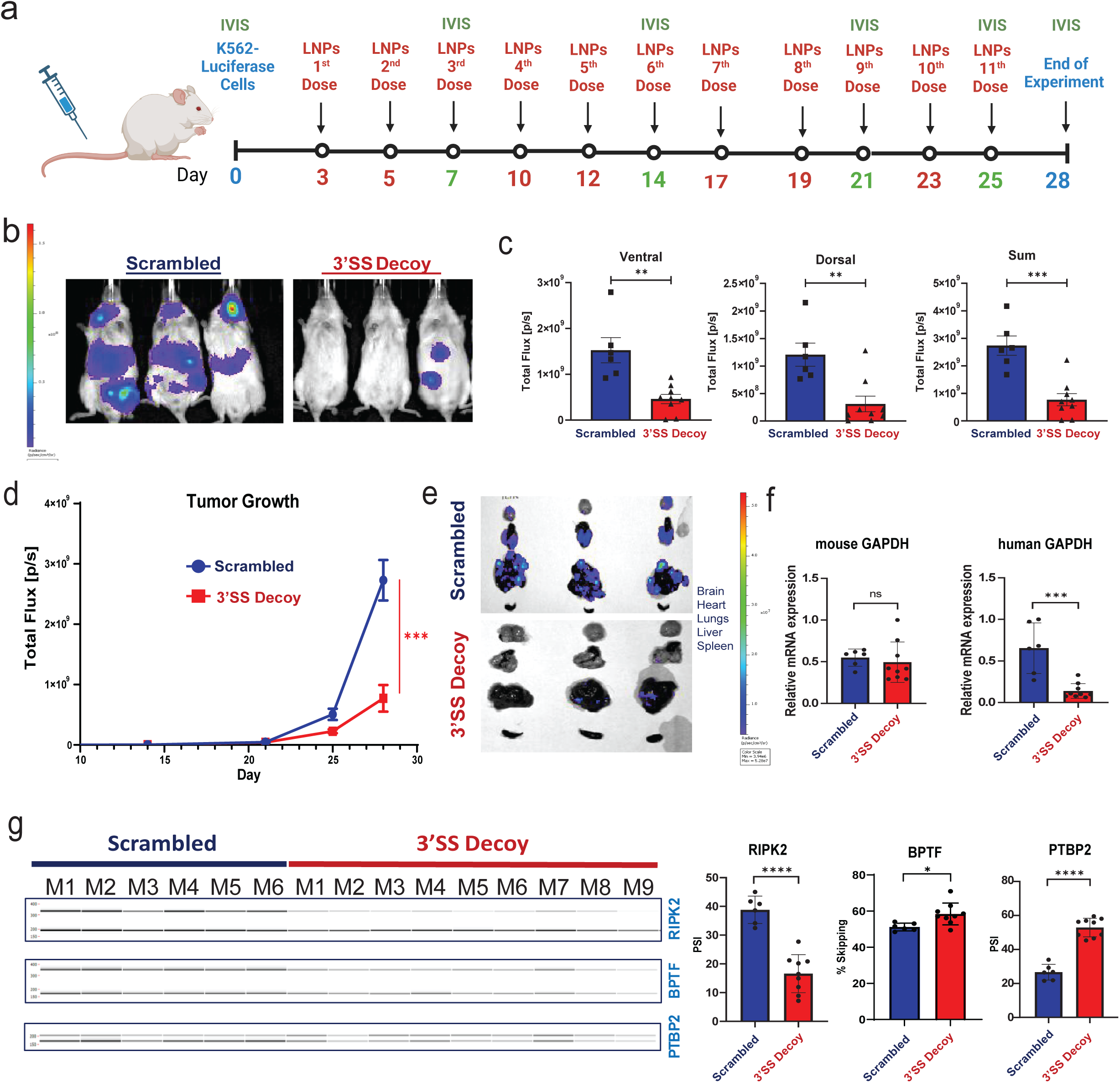
The 3’SS decoy decreases leukemia burden *in vivo*. (a) Scheme of the study design. On day 0, NSG mice were intravenously injected with 1 × 10^6^ luciferase-expressing K562 cells. Beginning on day 3, mice received intravenous administrations of LNP-encapsulated scrambled control or 3’SS decoy oligonucleotides (1mg/kg) every 2-3 days. Leukemia progression was monitored by weekly bioluminescence imaging (IVIS). Group sizes: scrambled (n = 7, one mouse died from the cancer on day 21) and 3’SS decoy (n = 9). (b) Representative ventral IVIS images at day 28 (endpoint) of three mice per group. Actual group sizes: scrambled (n = 7, one mouse died from the cancer on day 21) and 3’SS decoy (n = 9). (c) Quantification of total photon flux [p/s] at day 28, including ventral, dorsal and combined signals for each group. Unpaired two-tailed Student’s *t*-test was performed on data. Data is shown as mean ± s.d. P-values: ** p < 0.01, *** p < 0.001. (d) Longitudinal quantification of total photon flux [p/s] over the 28-day experimental period. Unpaired two-tailed Student’s *t*-test was performed on data (Day 28, end of experiment). Data is shown as mean ± s.d. P-values: ns= no asterisk, p > 0.05, *** p < 0.001. (e) Ex vivo IVIS imaging of harvested organs (brain, heart, lungs, liver and spleen) at day 28. Representative ventral IVIS images at day 28 (endpoint) of three mice organs per group. Actual group sizes: scrambled (n = 7, one mouse died from the cancer on day 21) and 3’SS decoy (n = 9). (f) Quantitative real-time PCR (qRT-PCR) analysis of RNA isolated from mouse livers, quantifying the levels of mouse and human GAPDH. Unpaired two-tailed Student’s *t*-test was performed on data. Data is shown as mean ± s.d. P-values: ns: p > 0.05, *** p < 0.001. (g) RT-PCR analysis of validated U2AF1 splicing targets (RIPK2, BPTF and PTBP2) in RNA isolated from mouse livers. Mouse numbers represented as M1-M9. Left: representative LabChip electropherograms. Right: corresponding quantification of RT–PCR isoform ratios. Unpaired two-tailed Student’s *t*-test was performed on data. Data is shown as mean ± s.d. P-values: * p < 0.05, **** p < 0.0001.

## Discussion

Recurrent mutations in spliceosomal genes have revealed a central role for RNA splicing dysregulation in the pathogenesis of myeloid malignancies^29–31^, yet these alterations also expose a paradoxical dependency on residual spliceosomal function^32^. Building on recent advances in therapeutics targeting aberrant RNA splicing in myeloid malignancies^33,34^, we exploited a splicing-associated vulnerability through the development of a synthetic RNA decoy that interferes with 3′ splice-site recognition, thereby functionally perturbing a central component of spliceosome activity. Our findings establish that decoy-mediated sequestration of U2AF1 and associated factors is sufficient to reprogram splicing, impair leukemic cell fitness, and suppress disease progression in vivo, providing a mechanistically distinct approach to targeting splicing factor dependencies.

Mechanistically, our data demonstrate that the 3’SS decoy engages U2AF1 in a sequence-dependent manner while simultaneously recruiting additional spliceosomal components (**Fig. 1**), consistent with the formation of stalled or non-productive splicing assemblies. The enrichment of canonical U2AF1-binding motifs among affected splice sites (**Fig. 2e**) indicates that the decoy effectively competes with endogenous pre-mRNA substrates. Notably, the decoy is not strictly specific to U2AF1, but rather engages a broader network of 3′ splice site-associated factors, including proteins recognizing the polypyrimidine tract. This feature, while potentially reducing molecular specificity, may in fact enhance therapeutic efficacy by amplifying disruption at a critical regulatory node of spliceosome function. Furthermore, a notable benefit of decoy-based strategies lies in their functional specificity. By engaging the RNA-binding domain of splicing factors, they selectively disrupt RNA recognition while leaving other roles, such as protein-protein interactions, largely unaffected. In contrast, siRNA-mediated gene silencing reduces overall U2AF1 protein levels, thereby perturbing not only its canonical splicing activity but likely also its non-canonical functions, including the recently identified roles in translation regulation^35^ and mitochondrial mRNA localization^36^, rather than selectively inhibiting its RNA-binding activity.

At the transcriptomic level, decoy treatment induces widespread and coordinated alterations in alternative splicing, accompanied by secondary transcriptional changes (**Fig. 2b, c**). Importantly, these splicing perturbations are not merely correlative but functionally consequential. By directly modulating individual U2AF1-regulated targets, including *RIPK2* and *BPTF*, we demonstrate that specific splicing events are sufficient to recapitulate key aspects of the decoy-induced phenotype (**Fig. 4**). These findings support a model in which cumulative disruption of multiple splicing-dependent pathways converges to impair leukemic cell survival, rather than reliance on a single dominant effector. The findings indicate that RIPK2 and BPTF function as downstream effectors of U2AF1-mediated splicing regulation and support the conclusion that the phenotypic consequences of decoy-mediated U2AF1 inhibition are, at least in part, mediated through these targets. This distributed vulnerability may represent a fundamental property of cancers driven by dysregulated RNA processing.

A major challenge in the development of RNA-based therapeutics is the efficient and selective delivery of oligonucleotides to target cells. Here, we show that LNP-mediated delivery enables robust intracellular delivery of the decoy in leukemic cells, including primary patient-derived samples, and is sufficient to drive functional and phenotypic effects (**Fig. 3 and Supplementary Fig. 3**). The ability of LNPs to facilitate systemic delivery and achieve therapeutic efficacy in vivo underscores their potential as a clinically viable platform for targeting RNA-protein interactions. While the observed preferential uptake in leukemic cells is encouraging, the mechanisms underlying this selectivity remain to be fully elucidated and may involve differences in membrane composition, metabolic state, or endocytic activity between malignant and normal hematopoietic cells.

Our in vivo studies further demonstrate that systemic administration of the decoy significantly reduces leukemic burden and induces splicing alterations consistent with on-target activity (**Fig. 5**), establishing proof-of-principle for therapeutic modulation of splicing through decoy-based strategies. These findings are particularly notable given the essential nature of the spliceosome, raising the possibility that partial and selective perturbation of splicing factor activity may be sufficient to achieve therapeutic benefit while avoiding overt toxicity. Nevertheless, a comprehensive evaluation of potential effects on normal hematopoiesis and other rapidly proliferating tissues will be critical for future translational development. Importantly, although it is known that LNPs significantly accumulate in the liver and spleen^37^, we did not detect major toxicity in treated mice (weight loss) (**Supplementary Fig. 4e**).

Several limitations warrant consideration. First, although U2AF1 serves as a primary molecular readout in this study, the decoy engages multiple components of the 3′ splice site recognition complex, and disentangling the relative contribution of each factor to the observed phenotype will require further investigation. Second, while our data supports a model of competitive sequestration, the precise structural and kinetic properties of decoy-protein interactions remain to be defined. Third, long-term effects, potential adaptive responses, and the impact of repeated dosing in vivo remain open questions that will be important for clinical translation. Fourth, extending this approach to additional disease models, including primary AML and MDS systems with diverse genetic backgrounds, will be essential to fully establish its therapeutic scope. Finally, this decoy oligonucleotide is expected to target both the wild type and mutant U2AF1, as it is not mutant-specific. Since mutations in splicing factors such as U2AF1 and SRSF2 are known to alter their sequence recognition preferences^38,39^, it will be interesting to determine whether engineering decoy oligonucleotides that mimic mutation-specific binding motifs can confer selective targeting of mutant splicing factors while sparing their wild-type counterparts.

Despite these challenges, our study provides a framework for targeting essential RNA-protein interactions using synthetic decoys. More broadly, the ability to pharmacologically mimic RNA elements and competitively engage protein binding interfaces represents an emerging paradigm with implications extending beyond splicing to multiple aspects of RNA biology.

In summary, we demonstrate that decoy-mediated targeting of 3′ splice site recognition disrupts spliceosomal function, rewires the leukemic transcriptome, and suppresses disease progression. These findings establish RNA decoy oligonucleotides as a mechanistically precise and therapeutically actionable approach to exploit splicing dependencies in cancer, and highlight the potential of LNP-based delivery systems to enable the clinical translation of RNA-targeted therapies.

**Supplementary Figure 1: Characterization and validation of the 3’SS decoy oligonucleotide.**

(a) Cytoplasmic and nuclear fractions were isolated from K562 cells and analyzed by immunoblotting for the indicated splicing factors; tubulin served as a cytoplasmic loading control.

(b) Quantification of proteins enriched by pull-down followed by mass spectrometry protein analysis using biotinylated 3’SS decoy or scrambled control oligonucleotides incubated with K562 nuclear extracts. Unpaired two-tailed Student’s *t*-test was performed on data. Data is shown as mean ± s.d. P-values: ns= no asterisk p > 0.05, * p < 0.05, ** p < 0.01, **** p < 0.0001.

(c) STRING analysis of the functional protein association network of significantly enriched proteins (n = 343; p < 0.05, FDR < 0.05). Protein clustering into three groups was performed using the k-means clustering algorithm.

(d) Gene Ontology enrichment analysis of biological processes associated with significantly enriched proteins identified by mass spectrometry (n = 343; p < 0.05, FDR < 0.05). The top 10 enriched pathways are shown.

In all panels, *n* ≥ 3 independent experiments.

**Supplementary Figure 2. The 3’SS decoy oligonucleotide induces leukemic cell sensitivity mediated through expression and splicing changes.**

(a) PCA based on normalized gene expression levels across samples. Each point represents a sample and is colored according to the experimental condition, as indicated in the legend. The PCA shows separation of samples according to treatment based on global gene expression profiles.

(b) Heatmap of differentially expressed genes based on shrunken log₂ fold-change (LFC) values, estimated by DESeq2. Rows represent genes and columns represent individual samples. Hierarchical clustering was applied to group genes with similar expression patterns across the different conditions.

(c) MA plot showing differential gene expression between 3’SS decoy-treated and scrambled control samples. Each point represents a gene. The x-axis indicates the mean of normalized counts, and the y-axis shows the shrunken log₂ fold-change estimated by DESeq2. Genes with p-value < 0.05 are highlighted in blue, while non-significant genes are shown in gray.

(d) Representative sashimi plots are shown illustrating RNA-seq read coverage and splice junction support for selected alternatively spliced events identified in the analysis. For each event, the plots display read coverage across the exon and its flanking regions, along with arcs indicating splice junction reads connecting adjacent exons. Samples are shown for untreated, scrambled control, and 3’SS decoy-treated conditions. These plots illustrate the differential exon inclusion or exclusion patterns associated with 3’SS decoy treatment. Data are presented as mean ± SD from *n* = 3 independent experiments. Statistical significance was assessed using an unpaired two-tailed Student’s *t*-test. Primers were designed to amplify regions flanking the alternatively spliced exons (see Supplementary Table 1). Shown are representative LabChip electropherograms and corresponding quantification of RT-PCR products from RNA isolated from K562 cells treated with the indicated scrambled control, 3’SS decoy oligonucleotide, control siRNA, siU2AF1- encapsulated LNPs at 500nM. Unpaired two-tailed Student’s *t*-test was performed on data. Data is shown as mean ± s.d. P-values: ns= no asterisk p > 0.05, * p < 0.05, ** p < 0.01, *** p < 0.001, **** p < 0.0001.

(e) Quantitative real-time PCR (qRT–PCR) analysis of RNA isolated from K562 cells treated for 72 h with LNP-encapsulated U2AF1-targeting siRNA. Relative U2AF1 mRNA expression was normalized to Actin; representative data are shown. Unpaired two-tailed Student’s *t*-test was performed on data. Data is shown as mean ± s.d. P-values: ns= no asterisk p > 0.05, **** p < 0.0001.

(f) Immunoblot analysis of protein lysates from K562 cells treated with LNP-encapsulated U2AF1-targeting siRNA for 72 h. Negative control siRNA and untreated cells were included as controls (left).

In all panels, *n* ≥ 3 independent experiments.

**Supplementary Figure 3: Efficient delivery of the 3’SS decoy into leukemic cells**

(a) Schematic representation of formulated LNPs encapsulating either a scrambled control or the 3’SS decoy oligonucleotide. Encapsulation efficiency measured by the RiboGreen assay (left). Particle size and polydispersity index (PDI) determined by dynamic light scattering (Zetasizer); *n* = 3 (middle and right).

(b) Quantification of Figure 2a. K562 cells were incubated for 24 hours at 37°C with Cy3-labeled RNA oligonucleotides (scrambled or 3’SS decoy) encapsulated in Cy5-labeled LNPs. Quantification of Cy3 and Cy5 fluorescence signals using a slide-scanner imaging system. Unpaired two-tailed Student’s *t*-test was performed on data. Data is shown as mean ± s.d. P-values: **** p < 0.0001.

(c) CellTrace Violet (Pacific Blue)- labeled peripheral blood- derived normal white blood cells (WBCS, green) were co-cultured with K562 cells (blue) at a ratio of ∼90% WBCs and ∼10% K562 cells and incubated with increasing concentrations (250ng, 500ng, 1000ng) of scrambled oligonucleotide encapsulated LNPs for 1 hour at 37°C. Flow cytometry analysis was performed, acquiring 10,000 events per sample.

(d) Quantification of flow cytometry plots in (c). Data is shown as mean ± s.d. P-values: *** p < 0.001.

(e) Peripheral blood cells from three leukemia patients were incubated with increasing concentrations (250ng, 500ng, 1000ng) of scrambled oligonucleotide encapsulated LNPs for 1 h at 37 °C, followed by flow cytometric quantification of Cy5 labeled LNP uptake. Flow cytometry analysis was performed with 10,000 events acquired per sample.

**Supplementary Figure 4: Additional experiment showing that the 3’SS decoy decreases leukemia burden in vivo**

(a) Schematic of the in vivo study design. On day 0, NSG mice were intravenously injected with 1 × 10^6^ luciferase-expressing K562 cells. Beginning on day 3, mice received intravenous administrations of lipid nanoparticle (LNP)- encapsulated scrambled control or 3’SS decoy oligonucleotides (1 mg/kg) every 2-3 days. Leukemia progression was monitored by weekly bioluminescence imaging (IVIS). Group sizes: scrambled (n = 4) and 3’SS decoy (n = 5).

(b) Representative ventral and dorsal IVIS images on days 18, 20, and 24 (endpoint).

(c) Longitudinal quantification of total photon flux [p/s] over the 28-day experimental period. Unpaired two-tailed Student’s *t*-test was performed on data. Data is shown as mean ± s.d. P-values: ns= no asterisk p > 0.05, * p < 0.05.

(d) Ex vivo IVIS imaging of harvested organs (brain, heart, lungs, liver and spleen) at day 24.

(e) Weight of the mice throughout the experiment described in Figure 5. Group sizes: scrambled (n = 7, one mouse died from the cancer on day 21) and 3’SS decoy (n = 9). Each line on the graph represents a mouse.

## Methods

### Cell Culture

K562 cell line was cultured in RPMI containing glutamine supplemented with 10% fetal bovine serum and 1% pen-strep. HEK-293T cells were cultured in DMEM containing glutamine supplemented with 10% fetal bovine serum and 1% pen-strep. To ensure the absence of mycoplasma contamination, all cell lines used in this study were routinely screened with the Vazyme MycoBlue Mycoplasma Detector Kit (cat # D101). Leukemia patient samples were obtained from Dr. Eran Zimran under written informed consent with the approval of the Hadassah Medical Center institutional review board.

### RNA Extraction, cDNA synthesis, RT-PCR

Total RNA was isolated using TRI Reagent (Sigma cat # T9424). For cDNA synthesis, 1 µg of total RNA was reverse transcribed to cDNA with iScript cDNA Synthesis kit (Bio-Rad cat #1708891). RT-PCR was conducted on 1 µl of cDNA using PCRBIO HS Taq Mix Red kit (BIOSYSTEMS cat# PB10.23.02) to confirm splicing and splicing modulation by the decoy oligonucleotides and the CRISPR-Cas9 system. PCR conditions were as described in the manufacturer’s protocol. PCR products were separated either on a 2% agarose gel or using LabChip analyzer.

### qRT-PCR

Total RNA was extracted with TRI Reagent (cat # T9424), and 1 μg of total RNA was reverse transcribed using iScript cDNA Synthesis kit (Bio-Rad, cat#1708891). qPCR was performed on the cDNA using iTaq Universal SYBR Green Supermix (Bio-Rad cat#1725124) and a Bio-Rad CFX384 Touch Real-Time System. Normalization was performed using HPRT or B-Actin primers. Primers are listed in **Supplementary Table 1**.

### Western blot analysis

Cells were lysed in Laemmli buffer and lysates were separated in 4-20% precast Mini-Protean TGX Stain-Free Gels (cat# 4568094) gels, then transferred to PVDF membranes using the Trans-Blot Turbo Transfer Pack (cat #1704157). The membranes were probed with primary and secondary antibodies listed in **Supplementary Table 2**. Quantification was performed using Image Lab 5.0 (Bio-Rad).

### Lentiviral infection and CRISPR-Cas9 directed splicing modulation

Lentiviruses were produced by co-transfection of HEK293T cells with psPax2 (Addgene #12260), pMD2.G (Addgene #12259) and CRISPR or pLVX Luciferase plasmids using FuGENE-HD (Promega cat #E2312) and OptiMEM (Gibco cat #51985-026). One day after transfection, the medium was replaced, and 48h after transfection, viruses were collected and filtered through a 0.45 µm membrane. K562 cells were infected with the viruses with the addition of polybrene (10 μg/ ml, Sigma cat# 107689). Selection with puromycin (2 μg/ ml, Sigma cat# P8833) was initiated 2 days after infection. Validation of the CRISPR-induced skipping event was performed by RT-PCR using specific primers for targeted splicing events.

For CRISPR-Cas9-directed splicing modulation, sgRNAs were designed to target the 3′ and 5′ end splice sites of the target exons in order to induce skipping of *RIPK2* and *BPTF*. The gRNAs were later cloned into LentiCRISPR v2 vector (Addgene #52961). A list of sgRNAs used in this study is provided in **Supplementary Table 3**.

### Lipid Nanoparticle (LNP) Preparation and Characterization

Ionizable lipid (DLin-MC3-DMA, Cayman Chemical, cat# 34364), cholesterol (Sigma, cat# C8667), 1,2-Distearoyl-sn-glycero-3-phosphocholine (DSPC) (Lipoid, cat# 556500), DMG-PEG2000 (Avanti, cat# 880151P), and DilC18(5)-DS (AAT Bioquest, cat# 22054) were mixed at a molar ratio of 50:38.5:10:1.5:0.036 with absolute ethanol in a tube. RNA payloads were suspended in 50 mM sodium acetate buffer (pH 4.5) (Sigma, cat# S7899). To create LNPs, the aqueous RNA phase was injected to the lipids EtOH phase using a 1mL tip under light vortex. The particles were then washed using Amicon Ultra-4 Centrifugal Filter unit (Millipore, cat# UFC10008) or Amicon Ultra-15 Centrifugal Filter unit (Millipore, cat# UFC10024) and were analyzed for size and uniformity by dynamic light scattering. Zeta potential was determined using the Malvern Zetasizer (Malvern, Worcestershire, UK). RNA encapsulation in LNPs was calculated according to the Quant-iT RiboGreen RNA assay kit (Thermo Fisher Scientific cat#R11491) by calculating the percentage encapsulation at 100% − (RNA-LNPs/RNA-LNPs with Triton X-100 (Sigma, cat#T9285).

### Pull down assay

K562 Nuclear Extracts (NE) were lysed in a buffer containing 20 mM Hepes, 1.5 mM MgCl_2_, 420 mM NaCl, 0.2 mM EDTA and 25% (v/v) glycerol. NE was incubated with 1μM biotin tagged oligonucleotides for 30 minutes at 30°C (see **Supplementary Table 4** for oligonucleotide sequences). Streptavidin beads (Thermo Scientific cat# 53117), previously blocked with a solution containing 50mg/ml heparin for 30 minutes at 4°C, were later added for 2 hours under rotation at 4°C. The pulled-down fraction was washed and then lysed in Laemmli buffer and analyzed by Western blot analysis.

### LC-MS/MS analysis

MS analysis was performed using a Q Exactive-HF mass spectrometer (Thermo Fisher Scientific) coupled on-line to an Ultimate 3000 Dionex (Thermo Fisher Scientific,) UHPLC. Peptides were separated on an acetonitrile gradient run at a flow rate of 150 nl/min from 4% to 50% over 57 minutes on a reverse phase 25-cm-long C18 column (Aurora Ultimate XT 25×75, ionopticks, AU). Survey scans (300–1,650 m/z, target value 3E6 charges, maximum ion injection time 20 ms) were acquired and followed by higher energy collisional dissociation (HCD) based fragmentation (normalized collision energy). A resolution of 60,000 was used for survey scans and up to 15 dynamically chosen most abundant precursor ions, with “peptide preferred” option selected, were fragmented (isolation window 1.6 m/z). The MS/MS scans were acquired at a resolution of 15,000 (target value 1E5 charges, maximum ion injection times 25 ms). Dynamic exclusion was 20 sec. Data were acquired using Xcalibur software (Thermo Scientific). To avoid carryover, the column was washed with 80% acetonitrile, 0.1% formic acid for and allowed to equilibrate in 1% acetonitrile between samples.

### MS data analysis

The data analysis was performed using MetaMorpheus version 1.0.5^40^. Carbamidomethyl on C was set as fixed modification and oxidation on M was set as variable. In addition, the GPTMD module was used for an open search of PTMs in the data. The data was searched against the human proteome database obtained from Uniprot and contained 104599 non-decoy protein entries including 296 contaminant sequences.

Statistical analysis was done using the Perseus software^41^. Prior to analysis contaminants and decoys were removed. Only protein groups with at least 3 valid intensity values were considered for further analysis. All remaining invalid values were imputed from normal distribution to allow for analysis.

Based on the significant protein names, the pathway enrichment analysis was conducted using the Reactome database^20^.

### Protein-protein interaction and network analysis (STRING)

Protein-protein interaction network analysis was performed using the STRING database^42^ to investigate functional associations among proteins significantly enriched in the mass spectrometry dataset. The list of differentially enriched proteins (n = 343; p < 0.05, FDR < 0.05) was uploaded to STRING and analyzed using the Homo sapiens background with default settings unless otherwise specified. Interaction networks were constructed based on known and predicted protein-protein associations. Functional clustering of the resulting network was performed within STRING using the k-means clustering algorithm, with the number of clusters set to three. The resulting clusters were visualized and used to identify major functional modules within the enriched protein set. Gene Ontology enrichment analysis of biological processes associated with the same significantly enriched proteins identified by mass spectrometry (n = 343; p < 0.05, FDR < 0.05) was performed. The top 10 enriched pathways are shown.

### Apoptosis assay

For decoy treated samples: K562 cells (0.5 × 10^6^ cells) were seeded on 12- well plates in triplicates. After seeding, the cells were treated with the different decoy oligonucleotides encapsulated in the LNPs. After 72h, cells were collected and apoptotic cell death was evaluated by using the Pacific Blue Annexin V Apotosis Detection Kit (BioLegend, cat# 640918) following the manufacturer’s instructions. For the CRISPR samples: after the termination of puromycin selection, cells were grown for 3 days in normal, selection-free, media and later collected and tested for cell death using the same kit.

### Cell-Titer Glo Viability Assay

To determine the cell viability following treatments, cells were treated with Cell Titer-Glo^®^ assay reagent according to the manufacturer’s instructions (Promega, cat# G7571). The obtained Relative Light Units (RLU) values corresponding to the amount of ATP present in the metabolically active cells were measured by a plate reader (BioRad).

### Human white blood cell purification

Collection of blood from patients was approved by the Hadassah Medical Center institutional review board. Following informed consent, blood samples (∼10cc) were collected from healthy volunteers. Blood samples were transferred to the lab for analysis no more than 15 minutes post-blood draw. Heparinized blood was mixed with an equal volume of Dextran 500 (3% in saline) and incubated 30 minutes at room temperature. The leukocyte-rich supernatant was collected and resuspended in 20ml 0.2% NaCl for 30 seconds to remove contaminating erythrocytes. Isotonicity was restored by the addition of 20 ml 1.6% NaCl. Cells were then washed three times in HBSS. Normal white blood cells were labeled with pacific blue using the Cell-Trace Violet Cell Proliferation Kit, cat# C34557A, and then co-cultured with K562 cells at different percentages, and incubated for 1 hour at 37°C.

### Flow Cytometry

Following the different incubations, cells were harvested and washed in FACS buffer (PBS supplemented with 0.5% BSA). Samples were analyzed on an BD LSR Fortessa Flow Cytometer (BD Biosciences), acquiring 10,000 events per sample. All experiments were performed with three independent biological replicates.

### Fluorescence Microscopy

After incubating K562 cells with the Cy5+ LNPs encapsulating Cy3+ oligonucleotides, the samples were collected, washed, fixed using 4% paraformaldehyde and cyto-centrifuged. Nuclei were stained with DAPI (VECTASHIELD Antifade Mounting Medium with DAPI, cat# H-1200). Images were taken on a Fluorescence Slide Scanner and processed using the NIS-Elements AR (4.40 version) software and analyzed on the QuPath software. Quantification was done on three independent biological samples, and each sample had three fields quantified.

### RNA-Seq Preprocessing and Alignment

Raw RNA-seq reads were processed using fastp (v0.23.2)^43^ to remove PCR duplicate reads and sequencing adapters. Quality control of each sample was performed using FastQC (v0.10.1)^44^, and results were summarized using MultiQC (v1.11)^45^. Processed reads were aligned to the human reference genome (GRCh38/hg38) and transcriptome annotation (GENCODE v28) using STAR (v2.7.3a)^46^.

RNA-seq data is generated as part of this study was deposited under the accession number <u>PRJNA1490051</u>: RNA sequencing of K562 cells treated with a 3’ splice site decoy oligonucleotide.

### Differential Gene Expression Analysis

Gene and transcript expression levels were quantified using Salmon (v1.4.0)^47^ Differential gene expression analysis between experimental conditions and their respective controls was performed using DESeq2 (v1.44.0)^48^. Genes were pre-filtered to retain those supported by at least 10 reads across samples. Log2 fold changes were estimated using the shrinkage LFC method. Genes were considered significantly differentially expressed if they met the following criteria: |log2 fold change| > 1 and adjusted p-value < 0.05.

### Alternative Splicing Analysis

Alternative splicing events were identified using PSI-Sigma (v2.3)^49^ applied to the aligned RNA-seq samples, using GENCODE v28 gene annotation as the reference transcriptome. The minimum number of supporting reads required for a splicing event was set to 10. Significant splicing events were defined as those with |ΔPSI| > 10%, p-value < 0.05, and FDR < 0.05.

PCA was performed based on the PSI values of detected splicing events using the prcomp function in R. Sashimi plots were generated using the *rmats2sashimiplot* package. For visualization purposes, intron lengths were scaled using the *--intron_s 10* parameter to improve readability of exon-intron structures. Based on the gene names, the pathway enrichment analysis was conducted using the Reactome database^20^.

### Sequence motif enrichment analysis

Sequence motif enrichment analysis was performed using XSTREME (MEME Suite v5.5.5)^50^. For each alternatively spliced region, sequences were extracted and extended by 250 bp upstream and downstream of the splice site.

Two analyses were performed. Motif enrichment was tested against a custom motif set representing known U2AF1 binding motifs: *UNAG, UUUUUUUUAG, UUNAG, and UUNAGCCC*.

Statistical significance of motif enrichment was evaluated using Fisher’s exact test or the binomial test, as implemented in the MEME suite^51^.

### Animal Studies

In vivo experiments were performed in accordance with the guidelines of IACUC at the Hebrew University. The study is in compliance with all the relevant ethical regulations. NSG (NOD.Cg- *Prkdc^scid^ Il2rg ^tm1Wjl^* /SzJ) mice were housed under standard laboratory conditions in specific-pathogen-free cages in an animal room at constant temperature (19–23 °C) and regulated humidity under a 12 h:12 h light-dark cycle and received standard laboratory chow and water ad libitum. All mice entered the experiments at ∼6 weeks of age. Both male and female mice were used for the experiments. In vivo imaging system (IVIS Spectrum) was used to monitor the growth of luciferase-labeled K562 cells; VivoGlo Luciferin, In Vivo Grade P1043, Promega cat#, was injected to the mice prior to imaging, according to the manufacturer’s protocol.

### Statistical Analysis

Tables and graphs for statistical analysis were created using GraphPad Prism 9 (GraphPad Software). *P* values < 0.05 were considered significant. Statistical significance between two groups was determined by the two-tailed Student’s *t*-test, and for experiments with more than two groups was determined by two-way ANOVA, details of statistical analyses are indicated in figure legends. All the data in the graphs are shown as mean ± s.d. unless stated otherwise. All experiments were performed a minimum of three independent times with similar results, each containing at least three technical replicates (wells) for each condition.

## Acknowledgments

This work was supported in part by grants from the Horizon Europe consortium grant (CANCERNA, 101057250), the Binational Science Foundation (BSF, 2021108), the Israel Science Foundation (ISF, 1601/23), and the Sasson Naor Estate Foundation.

Some figures were created with BioRender.com

