## Supplementary figures and images for "Targeting the 3’ splice site by a decoy oligonucleotide attenuates U2AF1 splicing activity and inhibits leukemia"

### Supplemental Figures

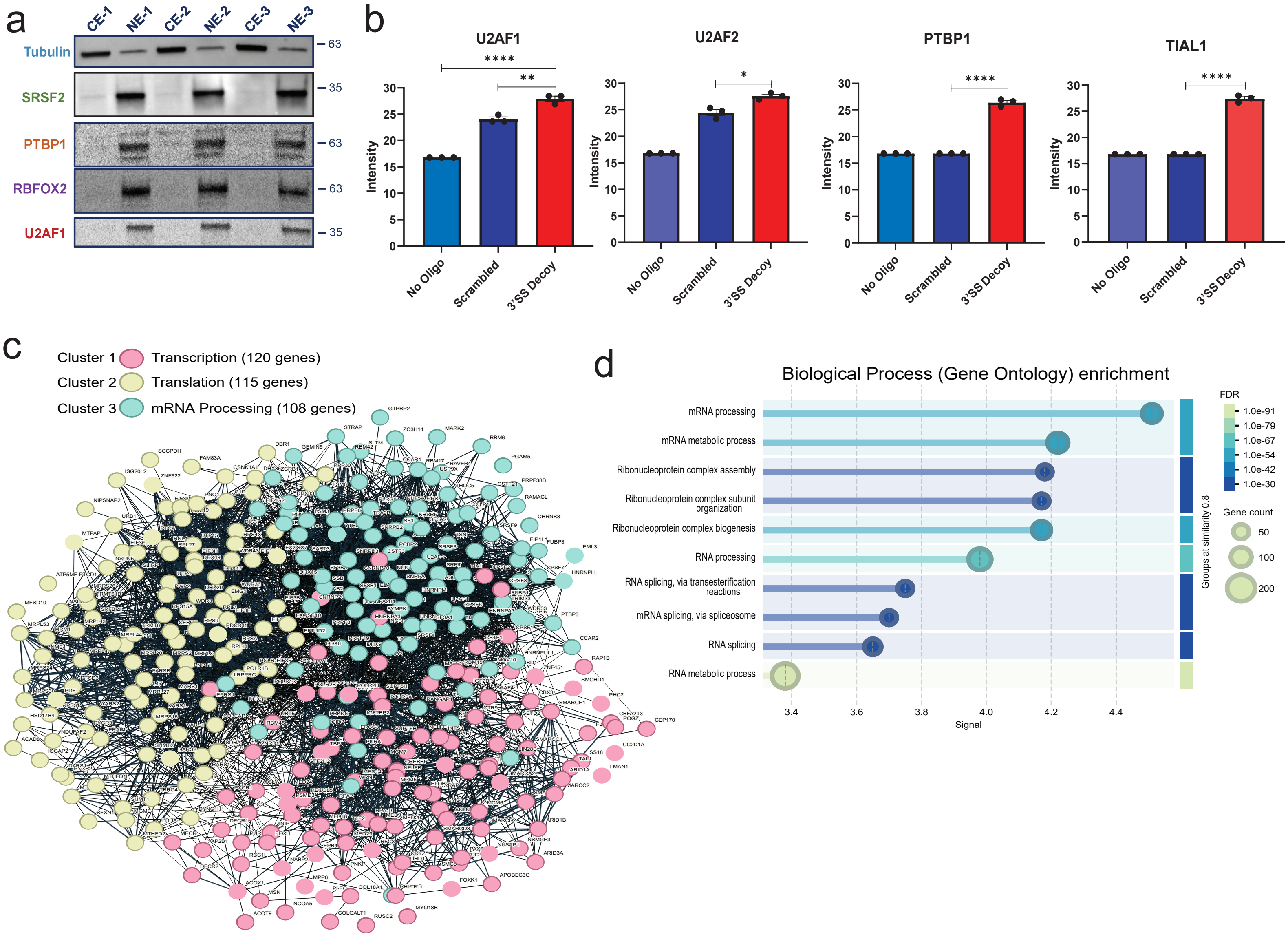


Supplementary Figure 1

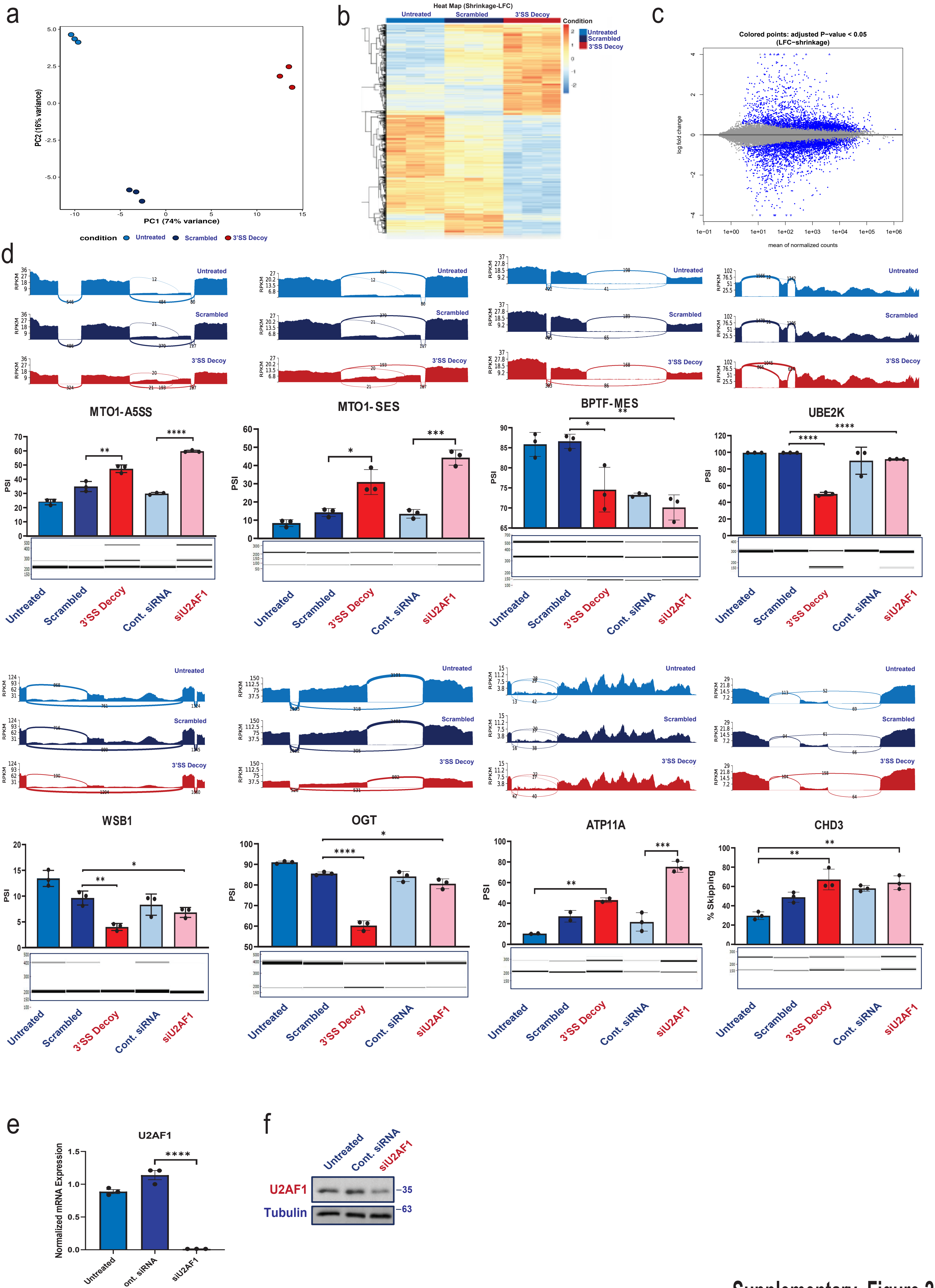


Supplementary Figure 2

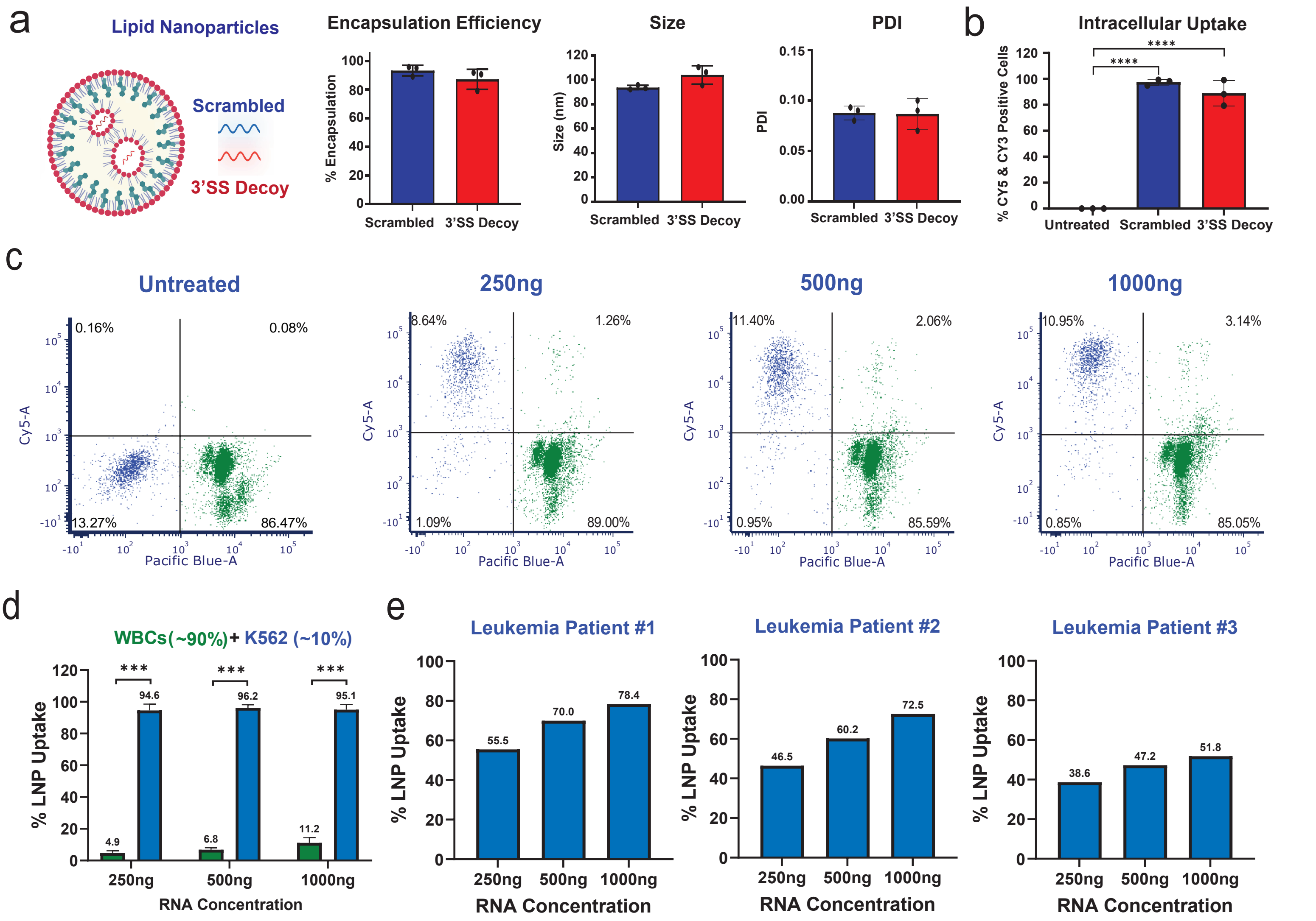


Supplementary Figure 3

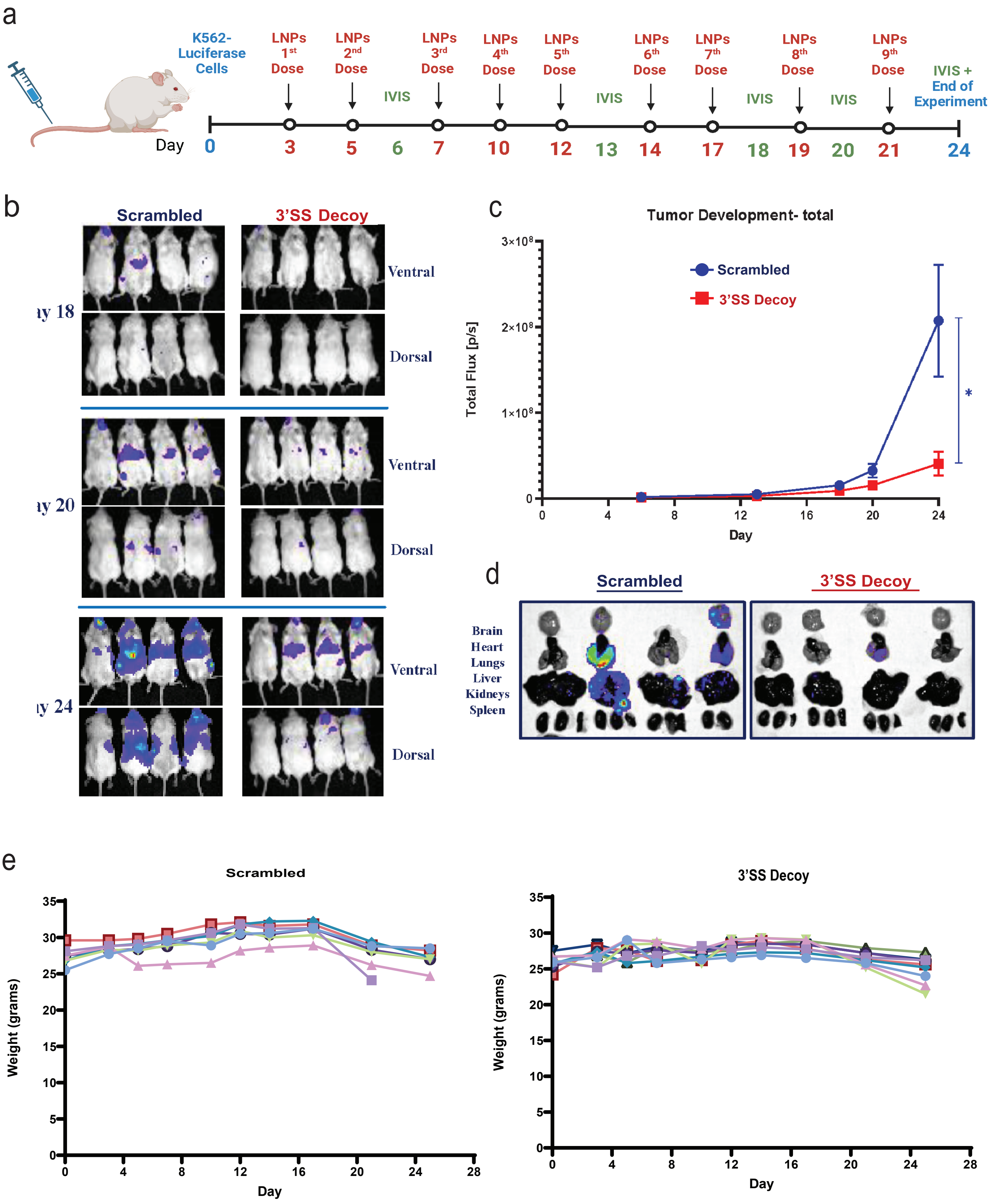


Supplementary Figure 4
