## Supplemental Tables for "Targeting the 3’ splice site by a decoy oligonucleotide attenuates U2AF1 splicing activity and inhibits leukemia"

**Supplementary Table 1: Primers**

| RT-PCR Primers | Forward | Reverse |
|---|---|---|
| RIPK2 (SES-EXON 2) | CATTCCCTACCACAAACTCG | AGGATGCGAAATCTCAATGG |
| ASUN (SES EXON 12) | TTGCACGTCCTTAGCAGTTC | TTAGGGCCCTTTCCTCTTGT |
| THYN1 (SES EXON 7) | AGAGGCTTACCCAGACCACA | CCTCCAGGCTCAAAACAAAA |
| PTBP2 (SES EXON 9) | AGCTGCTGCTGGCCGAGTG | GATTGGTTTCCATCAGCCATCTG |
| FLVCR1 (SES EXON 6) | TGCTGGAAGGATTGGGCTAA | TCAGGGTAAGTGATTTCAACAGC |
| BPTF (SES EXON 27) | AGGAAGCGGGAAGAGGAAAA | CGCCCATGGTACCAATTCTG |
| CRAMP1 (SES EXON 14) | GCTCCCTAGACCATCGGAAA | GTTCCAGTAGAGTCCTGCCC |
| APC (SES EXON 15) | TGTCTTGGCGAGCAGATGTA | TCCACAAAGTTCCACATGCA |
| MTO1 (SES EXON 7) | TTGCGTTTTCCAAACCGTCT | AAGGAAGGGGTGATCTGACG |

| | | |
|---|---|---|
| MTO1 (SES) | AAGGAAGGGGTGATCTGACG | 1: GCTCACCACAACCTCCCC<br><br>2: GGTGTTGGGCTTCATCTTCTG |
| BPTF (MES EXONS 5-6) | ACCAGATGATGACCCTGAGC | GTTTGCTATCTGGATTCCGCA |
| UBE2K (SES EXON 6) | CAGCTGCAATGACTCTCCG | TGTTGCAGTCTCTACATCCCA |
| WSB1 (A3SS EXON 6) | AATCTGACTATGACACTGAGCAT | 1: CGGAAACTAGAAGGACATCACC<br><br>2: CACCGGTCATTTGCTCCTC |
| OGT (SES EXON 7) | CCCTTGACCCAAACTTTCTGG | TGCCTCTTCAATGTTTCCCTG |
| ATP11A (SES EXON 29) | GCCGTTCCTCAACTACCAGA | GTGAATTCTGAGTCGCGGTC |
| CHD3 SES EXON 33) | CTAGAGGATGAGGTGCCAGG | CTGTCTTTTCCTCTCGTCGC |
| **qRT-PCR Primers** | | |
| B-Actin | GGCACCCAGCACAATGAAGA | AGGATGGAGCCGCCGATC |
| U2AF1 | GTGTACGTCAAGTTTCGCC | GCAGCAGGCTTCTCTGAAGT |
| Human GAPDH | GATCATCAGCAATGCCTCCT | TGTGGTCATGAGTCCTTCCA |
| Mouse GAPDH | ATCAAGAAGGTGGTGAAGCAG | CTTACTCCTTGGAGGCCATGT |

**Supplementary Table 2: Primary Antibodies**

| | |
|---|---|
| Tubulin | 1:5000, Abcam #ab6160, [YL1/2] |
| U2AF1 | 1:1000, Cell Signaling #13705S |
| SRSF2 | 1:1000, Thermo Scientific PA5-12402 |
| SRSF1 | 1:1000, (Caceres et al. 1997), mAb AK96 culture supernatant |
| SRSF6 | 1:1000, (Fu and Maniatis 1990), mAb 8-1-28 culture supernatant |
| PTBP1 | 1:10000, Abcam |
| HnRNPF/H | 1:1000, SC-32310, Santa Cruz Biotechnology |
| RBFOX2 | 1:1000, Sigma, #006240 |
| Peroxidase- conjugated AffiniPure Goat Anti-Mouse IgG (H+L) | 1:10000, Jackson ImmunoRese arch Inc., # 115- 035-003 |
| Peroxidase-conjugated AffiniPure Goat Anti-Rabbit IgG (H+L) | 1:10000, Jackson ImmunoRese arch Inc., # 111- 035-003 |

**Supplementary Table 3: guide RNAs**

| | |
|---|---|
| RIPK2 Guide 1 Forward | CACCGCATTAAATGAACTCCTACAT |
| RIPK2 Guide 1 Reverse | AAACATGTAGGAGTTCATTTAATGC |
| RIPK2 Guide 2 Forward | CACCGGTGTAATACTTACCCTATGT |
| RIPK2 Guide 2 Reverse | AAACACATAGGGTAAGTATTACACC |
| BPTF Guide 1 Forward | CACCGTGATTGTAAACGGGCACAAG |
| BPTF Guide 1 Reverse | AAACCTTGTGCCCGTTTACAATCAC |
| BPTF Guide 2 Forward | CACCGTGTTGGCATCACAGAAAAGG |
| BPTF Guide 2 Reverse | AAACCCTTTTCTGTGATGCCAACAC |

**Supplementary Table 4: RNA Oligonucleotides**

| Scrambled-GCGCGCX4 | Sequence (5' - 3') |
|---|---|

| | |
|---|---|
| Labeled either with CY3, Biotin, or none, at the 5' end. | eG*eC*eG*eC*oG*oC*oG*oC*oG*oC*oG*oC*oG*oC*oG*oC*oG*oC*oG*oC*eG*eC*eG*eC<br><br>Length: 24<br>RNA 2OMethyl: 16<br>MOE: 8<br>PTO: 23 |
| 3’SS Decoy<br><br>Labeled either with CY3, Biotin, or none, at the 5' end. | Sequence (5' - 3')<br>eT*eT*eT*eT*oU*oU*oU*oU*oA*oG*oC*oC*oC*oU*oU*oU*oU*oU*oU*oU*oU*oA*oG*oC*oC*oC*oU*oU*oU*oU*oU*oU*oU*oU*oA*eG*eC*eC*eC<br><br>Length: 39<br>RNA 2OMethyl: 31<br>MOE: 8<br>PTO: 38 |
| Control siRNA | SENSE: mC mG rU mU rA mA rUrCrG rCrGmU rAmUrA mArUmA rCrGrC rGrUmA T<br>ANTI SENSE: rArUmA rCrGrC rGrUmA rUmUrA mUrArC rGrCrG rArUrU rAmArC mGmAmC<br><br>Nomenclature: r= RNA, m= 2’O-Me |
| hs.Ri.U2AF1.13.1 | SENSE: mCmUrA mGrAmA rArGrU rGrUmU rGmUrA mGrUmU rGrArU rUrGmA C<br>ANTI SENSE: rGrUmC rArArU rCrAmA rCmUrA mCrArA rCrArC rUrUrU rCmUrA mGmCmA |

| | |
|---|---|
| | Nomenclature: r= RNA, m= 2’O-Me |
